# A super-resolution compatible workflow for highly multiplexed immunofluorescence of routinely processed kidney tissue

**DOI:** 10.1101/2023.12.11.570994

**Authors:** Florian Siegerist, Svenja Kitzel, Nihal Telli, Juan Saydou Dikou, Vedran Drenić, Christos E. Chadjichristos, Christos Chatziantoniou, Nicole Endlich

## Abstract

Deep insights into the complex cellular and molecular changes occurring during different (patho-)physiological conditions are essential for understanding the interactions and regulation of different proteins. This understanding is crucial for both research and diagnostics. However, the effectiveness of conventional immunofluorescence, an effective tool for visualizing the spatial distribution of cells or proteins, is limited in complex tissues. This is mainly due to challenges such as the spectral overlap of fluorophore wavelengths, a limited range of antibody types, and the inherent variability of samples.

Multiplex immunofluorescence imaging offers a solution to these limitations by enabling precise localization of proteins and identification of different cell types in a single tissue sample. In this study, we demonstrate the cyclic staining and de-staining of paraffin kidney sections, making it suitable for routine use and compatible with super-resolution microscopy for podocyte ultrastructural studies. We have further developed a computerized workflow for data processing which is accessible to all researchers through commercially available reagents and open-access image analysis codes.

As a proof of principle, we identified CDH2 as a marker for cellular lesions of sclerotic glomeruli in the nephrotoxic serum nephritis mouse model and cross-validated this finding with a human Nephroseq dataset indicating its translatability.

In summary, our work represents a significant advance in multiplex imaging, which is crucial for understanding the localization of numerous proteins in a single FFPE kidney section and the compatibility with super-resolution microscopy to study ultrastructural changes of podocytes.

## Introduction

The cellular heterogeneity of the kidney presents challenges when analyzing gene and protein expression. Although bulk omics can provide valuable insights, they have limitations in capturing the complete range of cell types involved in kidney (patho-)physiology. While still expensive and therefore inaccessible to a lot of researchers, single-cell RNA sequencing and spatial transcriptomics have emerged as powerful tools that enable the identification of individual cell types and their gene expression profiles in the kidney^1,2^. Immunofluorescence staining (IF) enables the sensitive localization of specific proteins in tissue sections. Besides this, IF has been used for gross glomerular morphometry^3^ or super-resolved ultra-morphometry of the filtration barrier^4^. Traditional IF is limited by spectral overlap of the fluorophores and the limited diversity of antibody species, which restricts the number of channels that can be imaged on one section. This limitation has been addressed by the highly multiplexed immunofluorescence (mIF) methods like the one developed by Gut and colleagues (4i), which uses cyclic staining and de-staining allowing the imaging of dozens of different proteins in cultured cells or tissue sections^5,6^. In the present study, we labeled proteins of interest with primary and secondary antibodies, which after being imaging with either confocal laser scanning microscopy (C-LSM) or super-resolution three-dimensional structured illumination microscopy (3D-SIM) are eluted from the sample (Fig. 1A). This way, the sole factor that limits the number of imaging cycles is the number of working antibodies available. To make this method as accessible as possible for a broad spectrum of researchers, all reagents used are commercially available, the image analysis codes are published open-source, and the imaging chambers are custom-designed and 3D-printed. Therefore, our approach opens new perspectives for inexpensive, rapid, and more efficient evaluation of histological samples not only in research, but also in clinical diagnosis because a limited amount of tissue can be used for multiple analyzes.

**Figure 1:**
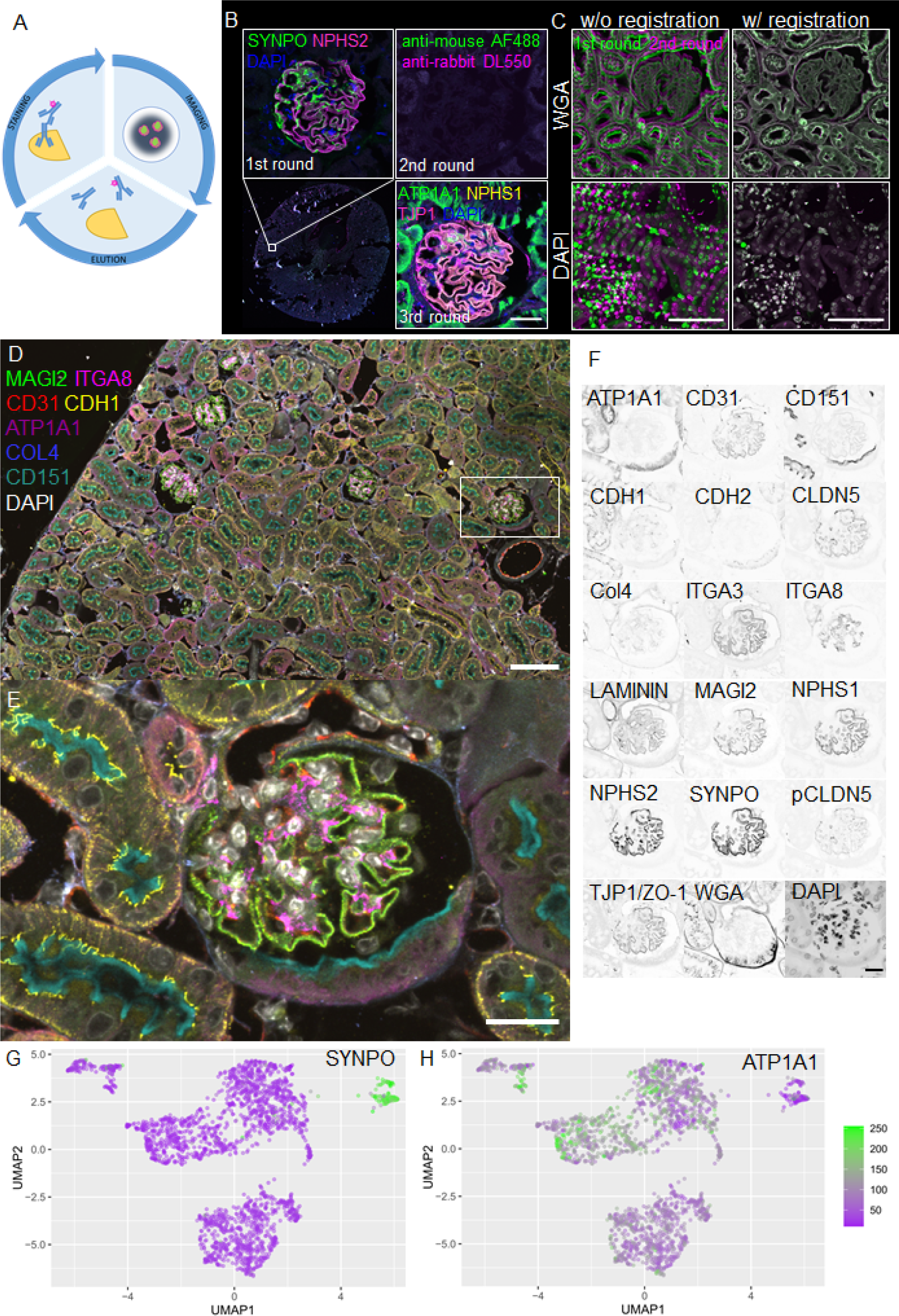
The scheme in A shows the general approach. A secondary immunofluorescence is performed, antibody binding is visualized with fluorescence microscopy, and the antibodies finally eluted from the section therefore enabling a new antibody binding and imaging cycle. The panel in B demonstrates the full elution of antibodies from a tissue section. A mouse kidney section was incubated with anti-synaptopodin (SYNPO) and anti-podocin (NPHS2) primary antibodies, which were detected with anti-mouse and anti-rabbit secondary antibodies (1^st^ round). After eluting the first secondary immunofluorescence, the section was incubated with the secondary antibodies only, which showed no binding of the secondary antibodies (2^nd^ round). Finally, the same section was again incubated with primary and secondary antibodies which demonstrated the integrity of the antigen on the section (3^rd^ round). The images in C show the functionality of the image registration algorithm to precisely align the consecutive imaging rounds. D, E and F show a merged mIF dataset for 17 different proteins, and DAPI as a nuclear marker of a mouse kidney section. Cells were segmented and single-cell protein mean fluorescence intensities were retrieved. A UMAP was calculated from the resulting dataset, demonstrating precise clustering according to diverse cell-identities as exemplarily shown for the upper right podocyte cluster (SYNPO as a marker protein: G) or tubular cells (ATP1A1: H). Scale bars represent 50 μm in B and C, and 20 μm in D-F.

## Short methods

### Tissue processing and immunostaining

Mouse kidney tissue was fixed with 4% paraformaldehyde (PFA) for a previous study has been used as described before^7^. Control rat tissue was fixed by retrograde aortic perfusion fixation with 4% PFA in deep anesthesia (Ketamin/Xylazin). This animal experiment has been approved and was overseen by the LALLF Rostock, Germany (file number 7221.3-1-037/22-1). After immersion fixation in 4% PFA overnight at room temperature, the tissue was embedded in paraffin and 2 μm sections were collected on 22x22mm #1.5 high-precision coverslips coated with 2% (3-Aminopropyl)triethoxysilane (Sigma-Aldrich, Cat. 440140). After deparaffinization, heat-induced epitope retrieval was performed by 5 min boiling in a pressure cooker in TRIS-EDTA buffer pH 9. Coverslips were immobilized in custom-designed and 3D-printed coverslip holders (.stl files available at http://www.github.org/Siegerist). Sections were blocked with 1% BSA, 200 mM NH_4_Cl, and 150 mM maleimide. Primary antibodies (diluted in 1% BSA; 200 mM NH_4_Cl) were incubated for 2 hours at room temperature (RT). Sections were washed with PBS, and secondary antibodies were incubated for 1 hour at RT. AlexaFluor 555 or CF770-conjugated wheat germ agglutinin (WGA) was added to the secondary antibody solution to a final concentration of 2 μg/ml. After several washes in PBS, cell nuclei were stained with 0.1 mg/ml DAPI. The sections were imaged in 700 mM N-acetyl-cysteine in H_2_O pH 7.4 on a FluoView3000 confocal laser scanning system (Olympus) equipped with a 20x 0.8 NA dry objective (UPLXAPO20X) or a UPLSAPO60XW NA 1.2 water immersion objective (validation experiments). After imaging, antibodies were eluted with 0.5 M L-glycine, 3 M urea, 3 M GC; 70 mM TCEP in ddH_2_O (pH 2.5), blocked, and re-stained as described above. Imaging with a three-dimensional (3D) super-resolution microscopy was performed by using a Nikon N-SIM E system (Nikon Instruments, Japan) at NIPOKA GmbH (Greifswald, Germany). For overview images the 20x/0.75NA objective was used and the high-resolution images were acquired by using 100x/1.35NA silicon immersion objective. The Nikon N-SIM E system is equipped with 488, 561 and 640 nm laser lines.

### Quantitative image analysis

A script was established in the IJ1 language that registers and corrects the drift which can be found at http://www.github.org/Siegerist. Briefly, minima and maxima are detected with sub-pixel accuracy in both respective images and translated to one another using a descriptor-based registration algorithm^8^. This process was iteratively repeated until all images were merged. From the overviews glomeruli were detected with a pretrained UNet, and individual cell nuclei with surrounding cytoplasm were segmented with a custom-trained Deep Learning network (StarDist in QuPath v.0.4.3)^9^. UMAPs were generated in R-Studio with the UMAP package.

## Results

### Optimization of the 4i protocol for FFPE kidney tissue

For cyclic staining, imaging, and de-staining (Fig. 1A) of kidney, we improved tissue adhesion by coating coverslips with 3-aminopropyltriethoxysilane which was superior to poly-l-lysine or uncoated slides (Suppl. Fig 1). During elution, both secondary and primary antibodies were efficiently removed from their epitopes as shown in Figure 1B.

As exemplarily shown for mouse and human kidney tissue in Suppl. Figure 2 and 3, in which the same glomerulus was imaged over 3 cycles, the staining, and de-staining worked efficiently without bleed-through from previous rounds and the tissue morphology was unimpaired.

To establish an mIF antibody panel, we screened 44 antibodies on human and mouse kidney tissue in single immunostainings (**Suppl. Table 1**). We found 20 antibodies for mouse kidney tissue including tubular markers (ATP1A1, E-cadherin, N-cadherin, ezrin), glomerular markers (podocin, synaptopodin, nephrin, MAGI2, ITGA3, DACH1, CD151), tight junction proteins (TJP1, CLDN5, phospho-CLDN5), endothelial markers (CD31) mesangial makers (ITGA8), and markers for extra cellular matrix (ECM) components (laminin, collagen IV) (Suppl. Fig. 4).

To reduce drift, staining and imaging cycles were performed in a custom 3D-printed chamber. As drift could not be mechanically abrogated completely (Fig. 1C), a marker that is consistently stained in every round is required to detect the shift. Since DAPI staining quality was inconsistent, independent on the number of staining cycles (Suppl. Fig. 5), we stained the surface using near-infrared CF770-conjugated wheat-germ agglutinin which could be efficiently eluted and re-stained consistently on the same section (Suppl. Fig. 6).

2D-descriptor-based image registration within the open-source ImageJ ecosystem was used to precisely align the datasets of two subsequent imaging rounds in both DAPI and WGA channels (Fig. 1C, overviews in Suppl. Fig. 7).

Large tile scans of tissue sections were acquired on a confocal laser scanning system and stitched to high-resolution overviews. To retrieve single-cell fluorescence data for all antigens, we established a workflow that preprocesses the raw images to generate precisely aligned multichannel data (Suppl. Fig. 8A).

Subsequently, the section was virtually dissected into glomeruli with a custom-trained U-Net and in single cells with a StarDist network (Suppl. Fig. 8B). For every cell segmented on the slide, morphometric measures (nuclear size, roundness, and nucleus-cytoplasm ratio), classification to a tissue compartment (glomerulus vs. tubulointerstitium), and fluorescence intensity quantifications were reported. In this way, single-cell co-expression analysis from a few thousand cells was enabled, as demonstrated for NPHS2, SYNPO, ATP1A1, and TJP1 with NPHS1, respectively (Suppl. Fig. 8C). For a typical overview of a tissue section of 700x700 μm as shown in Figure 1D, we could retrieve single-cell data from at least 3000 cells.

As shown in the magnified view in Figure 1E, the resolution and orientation of the cyclically acquired images was found to be sufficient to capture the details of the subcellular localization of the proteins studied. Notable examples are the distinct localization of CD151 at the brush border, ATP1A1 to the basolateral cell membrane, and collagen IV to the basement membrane. All three glomerular cell types could be identified by their staining such as MAGI2 (podocytes), ITGA8 (mesangial cells), and CD31 (glomerular endothelial cells), respectively. Figure 1F shows the single-channel overview of all targets stained on this tissue section. The image analysis workflow described above generated a highly multidimensional dataset that was used to cluster cells in an unbiased manner according to their expression patterns and morphometric measures using Uniform Manifold Approximation and Projection (UMAP). As shown in Figure 1 G and H, four well-separated clusters could be identified. The smallest cluster in the upper right could be identified as the podocyte cluster by the synaptopodin expression (SYNPO, Fig. 1G). The larger cell clusters represented tubular cells (ATP1A1, Fig. 1H).

### Applying mIF in a crescentic glomerulonephritis model identified CDH2 as a marker for cellular glomerular crescents

To benchmark our workflow, we used the antibody panel in murine nephrotoxic serum nephritis (NTS), a model for crescentic glomerulonephritis. The kidney of the NTS-injected animal showed glomerular hypertrophy (Fig. 2B) as well as a loss of podocytes (depletion of triple NPHS1-, NPHS2-, and MAGI2-positive intraglomerular cells). In glomerular cross-sections, we quantified 1.2 podocytes/1000 μm^2^ in NTS animals versus 3.9 podocytes/1000 μm^2^ in control animals (p<0.0001; Fig. 2C). Furthermore, a significant intraglomerular accumulation of collagen IV was detected (Fig. 2D). The proximal tubule marker CDH2 (N-cadherin) was found to be expressed particularly in cellular lesions of sclerotic glomeruli (0.6 CDH2+ cells/1000 μm^2^ in the NTS tissue (p=0.0464) versus no positive cell was found in the control animal (Fig. 2E). This finding was verified with standard immunofluorescence for CDH2 on an extended cohort of 6 animals (Fig. 2F) as well as in FSGS patients. In isolated glomeruli of FSGS patients, 5.4-fold upregulation of the CDH2 mRNA in contrast to healthy kidneys was seen (Fig. 2G, Hodgin FSGS Glom dataset in Nephroseq).

**Figure 2:**
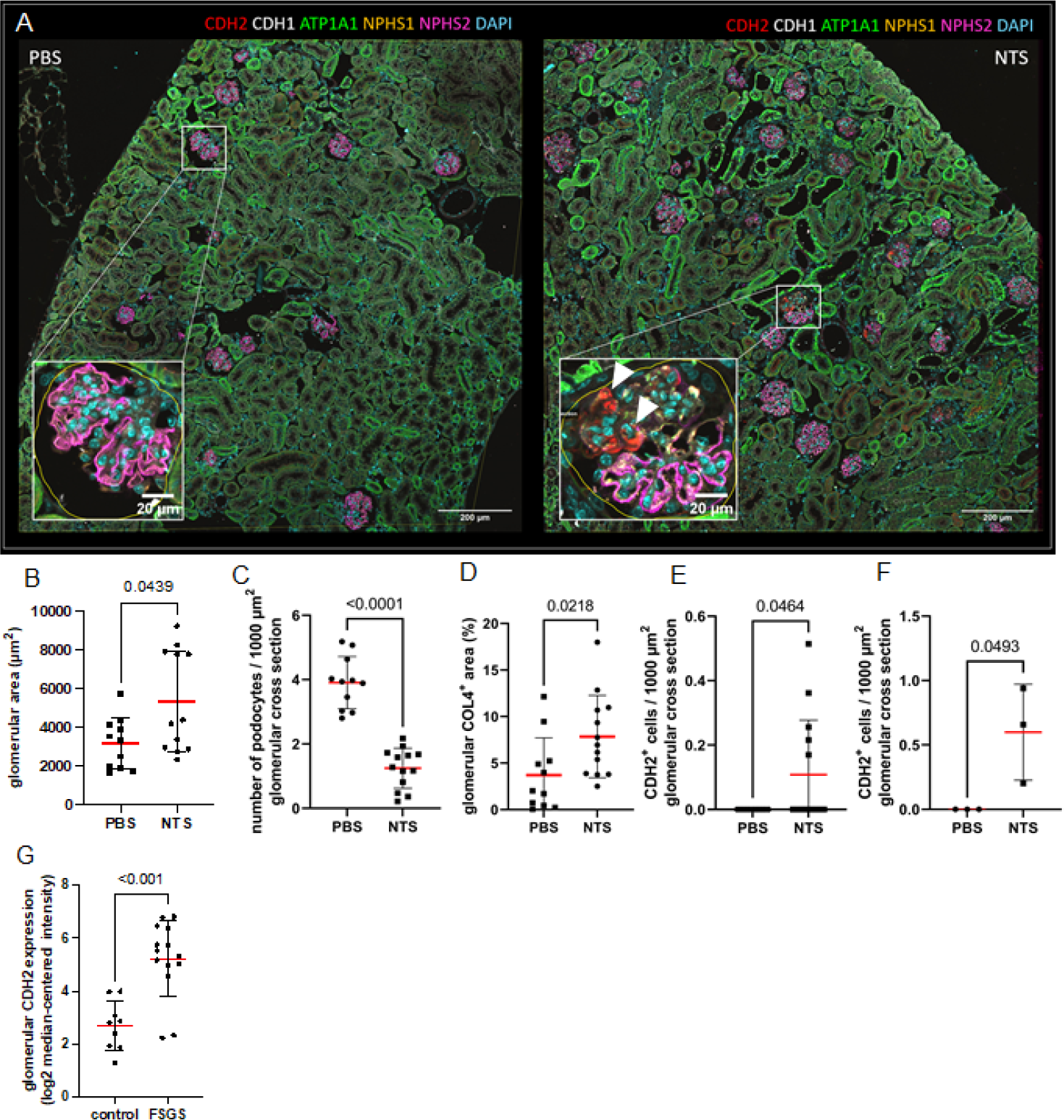
Figure 2A shows a comparison of a control mouse section (PBS) with an NTS-injected mouse. We demonstrate glomerular hypertrophy as shown by increase of the mean glomerular cross-sectional area (B). Additionally, decreased podocyte density in the NTS-injected animals (C) was demonstrated indicating podocyte depletion. Interestingly, we found aincrease of CDH2-expressing glomerular cells only in NTS-injected animals (arrowheads in A), which could be quantified as shown in D. To verify this finding, the analysis was extended to 3 animals per group in which the appearance of CDH2+ cells in the glomerular tuft was consistent (E). Within the Nephroseq database, CDH2 was enriched in glomeruli of patients diagnosed for FSGS (F).

### Super-resolved mIF uncovers glomerular filtration barrier details

To reveal the subtle morphological details of the filtration barrier, in particular the podocyte foot processes, and the filtration slit, we analyzed rat kidney sections using 3D structured illumination microscopy (3D-SIM). The tissue preservation allowed resolution of the filtration slit by staining for podocin, even after multiple previous staining, imaging, and elution cycles. As shown in the overview in Fig. 3A, we could super-resolve both glomerular (nephrin, integrin alpha 8, SSeCKS) and tubular marker proteins (ATP1A1, LRP2), as well as ubiquitously expressed ECM proteins (laminin). As demonstrated in the magnifications in Fig. 3B and C, the resolution was high enough to distinguish individual podocyte foot processes, therefore enabling podocyte foot process morphometry. These results demonstrate the direct integration of super resolution microscopy into the mIF workflow.

**Figure 3.**
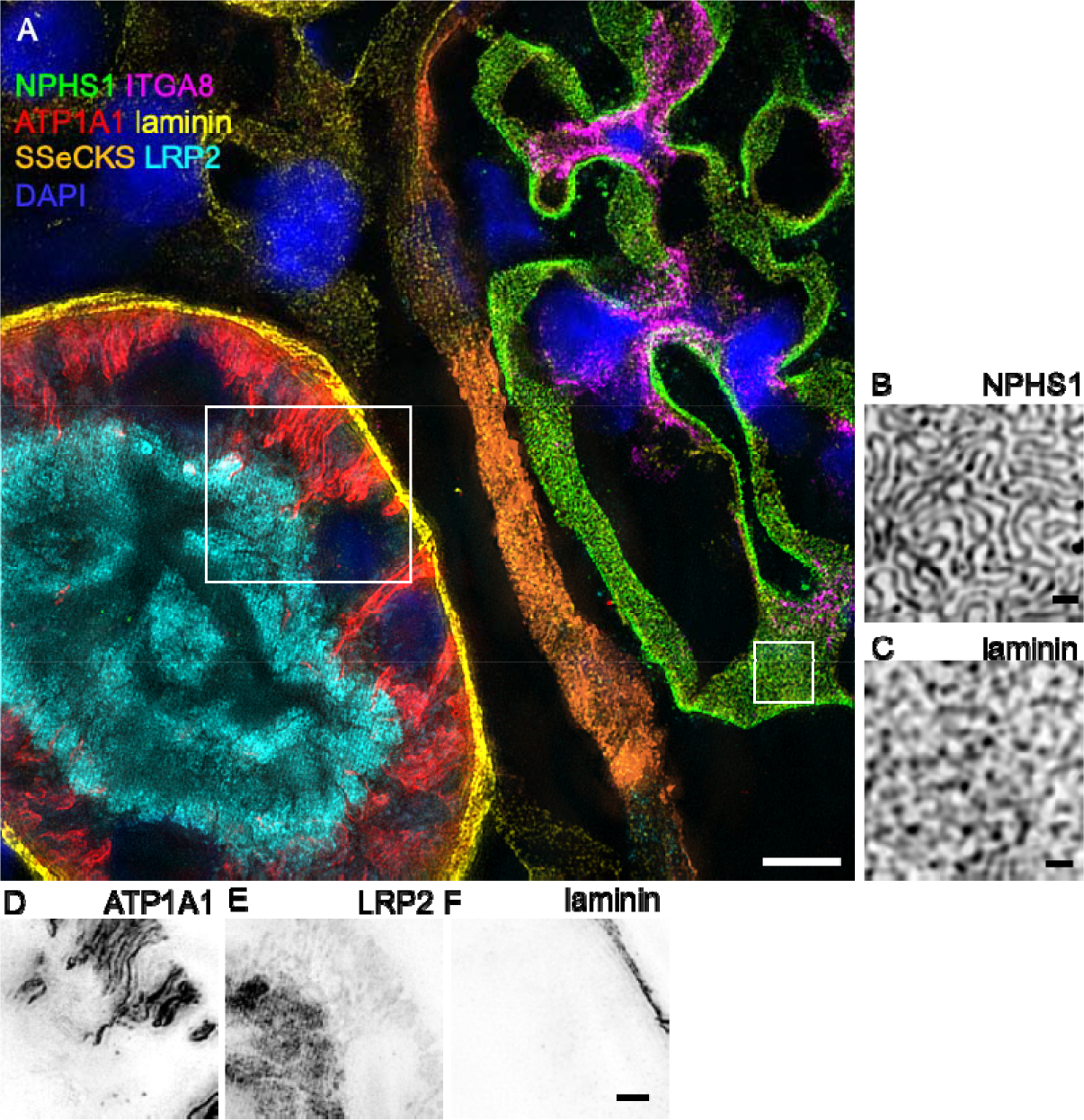
The image in A shows a super-resolved mIF dataset consisting of 6 different tubular and glomerular markers. Magnifications in B show that the resolution was high enough to determine podocyte foot processes on the laminin positive GBM (C). Besides glomerular proteins, also tubular markers like ATP1A1 in the basolateral membrane (D) and LRP2 in the apical cell membrane (E), as well as the tubular basement membrane (F) could be precisely resolved from one another. Scale bars represent 5 μm in A, 500 nm in B and C, and 2 μm in D-F.

## Discussion

Direct single-cell correlation of multiple proteins of interest is typically limited in 3-4-plex classic IF due to the problem of a limited number of different species for antibody production. However multiplex immunofluorescence analysis has the potential to significantly increase the amount of information about protein expression, localization as well as the relation to other proteins in single kidney section and therefore give a more comprehensive view of the tissue microenvironment. Until today, there are different techniques published which allow a multiplex staining of cells and tissue. Of the techniques available, the one presented by Gut et al. works with an adaption of routine secondary immunofluorescence utilizing commercially available antibodies and fluorophores^6^. Therefore, the barriers to establish this method in a multitude of labs are relatively low. Herein, we used only commercially available antibodies, so that the protocol presented here can be easily reproduced. While we used mIF with C-LSM and 3D-SIM, other fluorescence microscopic methods like wide field microscopy work with this approach as well.

Within the kidney field, also other mIF approaches have been published, like the one presented by Rajagopalan et al.^5^. However, this specific technique requires the direct conjugation of antibodies with modified nucleic acids, therefore increasing hands-on time, complexity, and costs of an mIF experiment.

One important advantage of our established method is that all reagents are commercially available and commonly used in classic IF, making it easy to extend existing workflows. Taken together, we present a straightforward addition to massively expand the information obtained from a single FFPE section with reagents available in most laboratories already working with IF.

Besides the establishment of the protocol for FFPE sections, we found CDH2 as a marker upregulated in cellular crescents of murine NTS-nephritis. While in healthy animals, CDH2 is localized in in the apical junctional complex of proximal tubule cells, its distribution changed in glomeruli of NTS-injected animals. Similarly it has been described by other groups and us, that cells at the interface between parietal epithelial and proximal tubular cells can be activated and recruited to injured glomeruli so that tubular markers are *de-novo* expressed in injured glomeruli^10,11^. Interestingly, this finding was consistent with a microarray dataset of laser-captured glomeruli from archived FSGS patients^12^. Therefore, following studies will focus on the establishment of CDH2 as a marker protein for the classification of glomerular injury in patients.

## Contributions

FS and NE conceptualized the study. FS and SK established the methodology. FS, SK, NT, and VD performed experiments and generated imaging data. FS and JSD established the quantitative image analysis workflow and trained DL networks. CC and CC contributed the NTS model to this study. FS, SK, NE wrote the manuscript and generated figures. All authors approved the final version of the manuscript.

## Acknowledgements

This work was supported by a grant of the Gerhard-Domagk Masterclass of the University Medicine Greifswald to FS and by a grant of the Federal Ministry of Education and Research (BMBF, grant 01GM1518B, STOP-FSGS) to NE. This work was generously supported by the Südmeyer fund for kidney and vascular research (‘Südmeyer Stiftung für Nieren-und Gefäßforschung’) and the Dr. Gerhard Büchtemann fund, Hamburg, Germany. CC and CEC were supported from the Inserm and SU annual recurrent funding.

## References

1. Park J, Shrestha R, Qiu C, et al. Single-cell transcriptomics of the mouse kidney reveals potential cellular targets of kidney disease HHS Public Access. Science (80-) 2018;360(6390):758–63.

2. Chung JJ, Goldstein L, Chen YJJ, et al. Single-cell transcriptome profiling of the kidney glomerulus identifies key cell types and reactions to injury. J Am Soc Nephrol 2020;31(10):2341–54.

3. Zimmermann M, Klaus M, Wong MN, et al. Deep learning-based molecular morphometrics for kidney biopsies. JCI Insight 2021;6(7).

4. Siegerist F, Ribback S, Dombrowski F, et al. Structured illumination microscopy and automatized image processing as a rapid diagnostic tool for podocyte effacement. Sci Rep 2017;7(1):11473.

5. Rajagopalan A, Venkatesh I, Aslam R, et al. SeqStain is an efficient method for multiplexed, spatialomic profiling of human and murine tissues. Cell reports methods 2021;1(2).

6. Gut G, Herrmann MD, Pelkmans L. Multiplexed protein maps link subcellular organization to cellular states. Science 2018;361(6401).

7. Artelt N, Ludwig TA, Rogge H, et al. The role of palladin in podocytes. J Am Soc Nephrol 2018;29(6):1662–78.

8. Hörl D, Rojas Rusak F, Preusser F, et al. BigStitcher: reconstructing high-resolution image datasets of cleared and expanded samples. Nat Methods 2019;16(9):870–4.

9. Siegerist F, Hay E, Dikou JS, et al. ScoMorphoFISH: A deep learning enabled toolbox for single-cell single-mRNA quantification and correlative (ultra-)morphometry. J Cell Mol Med 2022;26(12):3513–26.

10. Kuppe C, Leuchtle K, Wagner A, et al. Novel parietal epithelial cell subpopulations contribute to focal segmental glomerulosclerosis and glomerular tip lesions. Kidney Int 2019;96(1):80–93.

11. Hansen KUI, Siegerist F, Daniel S, et al. Prolonged podocyte depletion in larval zebrafish resembles mammalian focal and segmental glomerulosclerosis. FASEB J 2020;34(12):15961–74.

12. Hodgin JB, Borczuk AC, Nasr SH, et al. A molecular profile of focal segmental glomerulosclerosis from formalin-fixed, paraffin-embedded tissue. Am J Pathol 2010;177(4):1674–86.

