## Supplementary material for "A super-resolution compatible workflow for highly multiplexed immunofluorescence of routinely processed kidney tissue": Suppl.

*Corresponding address:

### Supplementary Figures

Suppl. Fig. 1

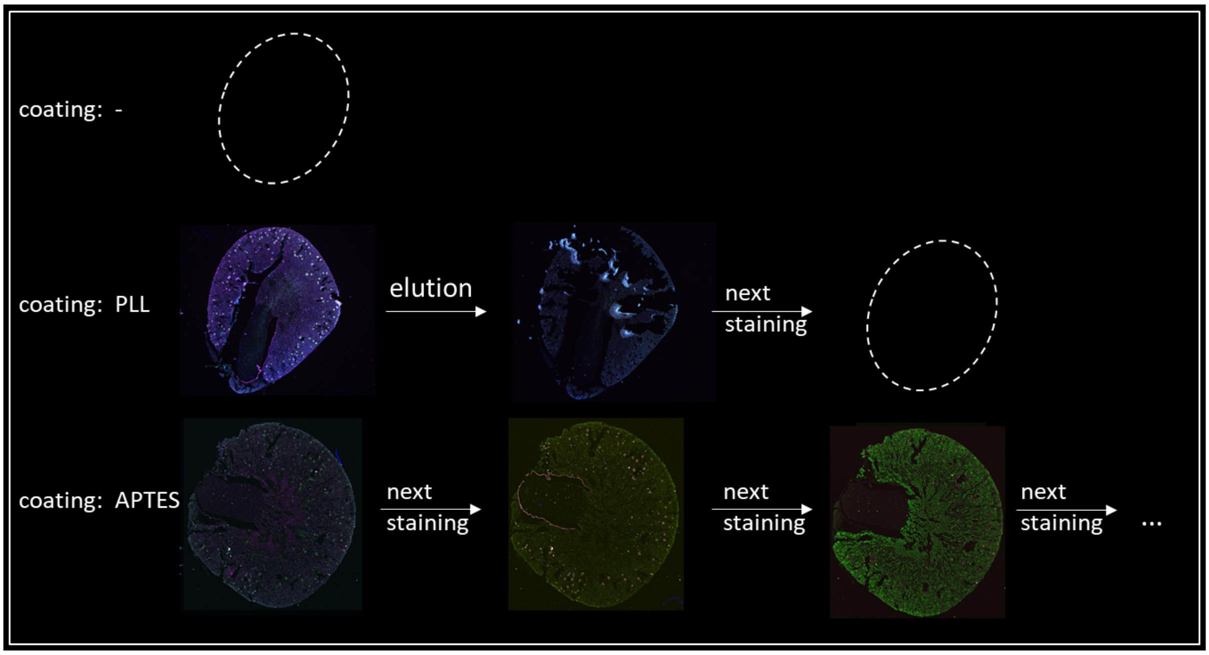


Suppl. Fig. 1 demonstrates the improved adherence of kidney tissue sections to class coverslip surface by APTES. Only after APTES coating, tissue remained on the slide during multiple staining, imaging, and elution cycles.

Suppl. Fig. 2

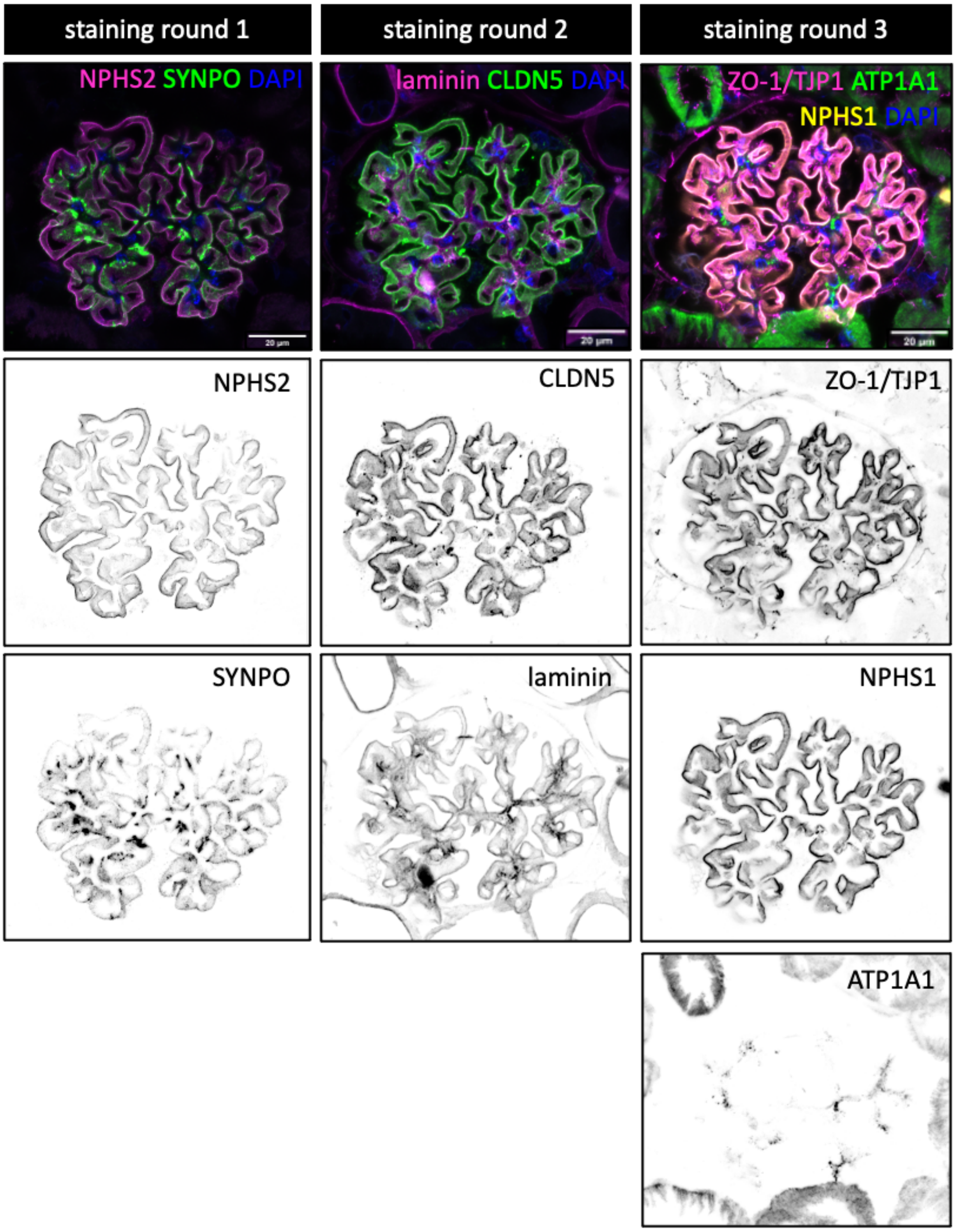


Suppl. Fig. 2: A single mouse glomerulus is shown over three consecutive staining and imaging rounds. A total of 7 marker proteins are imaged over the three rounds. No cross-bleed between individual staining rounds can be seen. The scale bars indicate 20 µm.

Suppl. Fig. 3

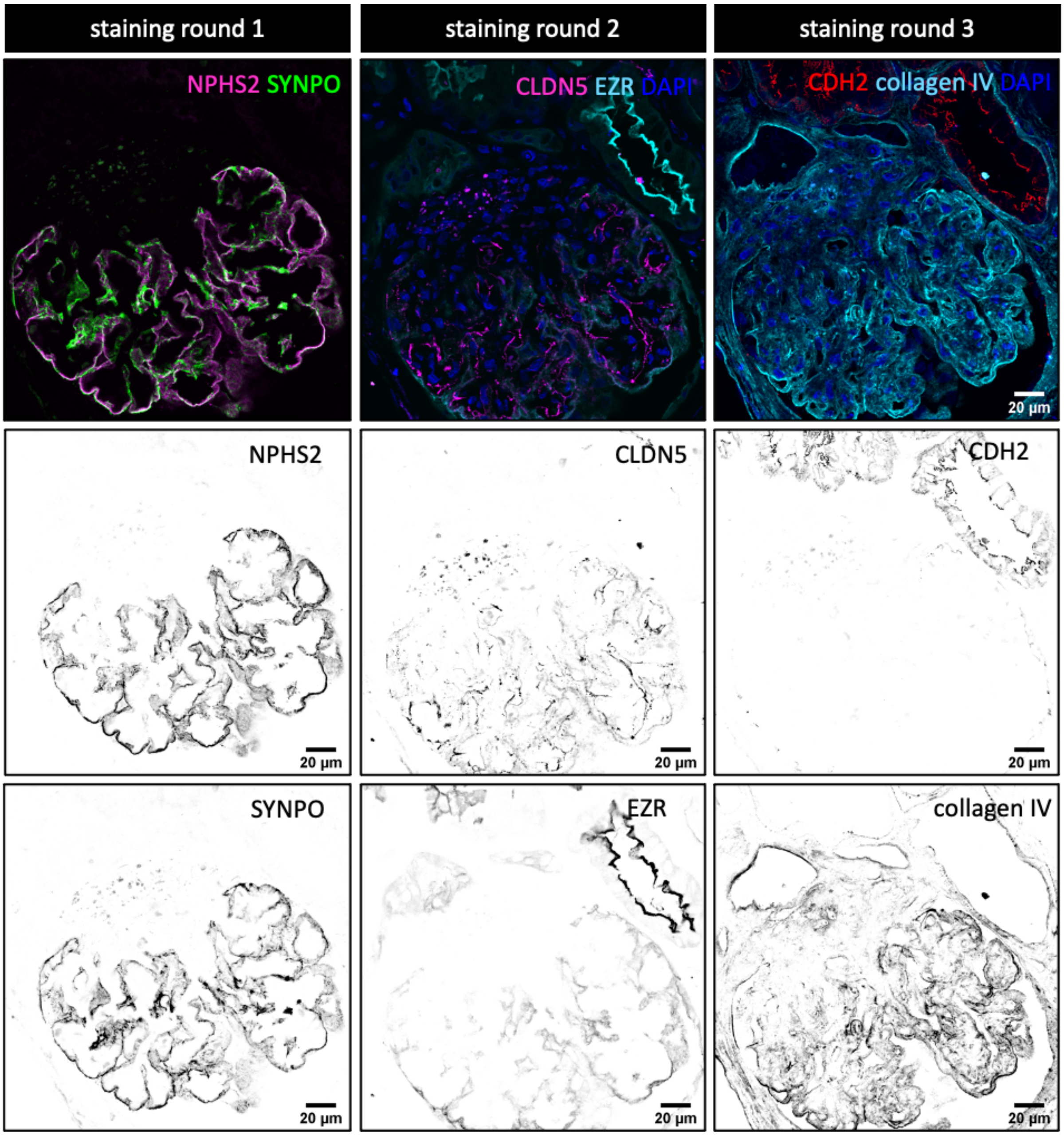


Suppl. Fig. 3: A single human glomerulus from an FFPE section is shown over three consecutive staining and imaging rounds. A total of 6 marker proteins are imaged over the three rounds. No cross-bleed between individual staining rounds can be seen. The scale bars indicate 20 µm.

Suppl. Fig. 4

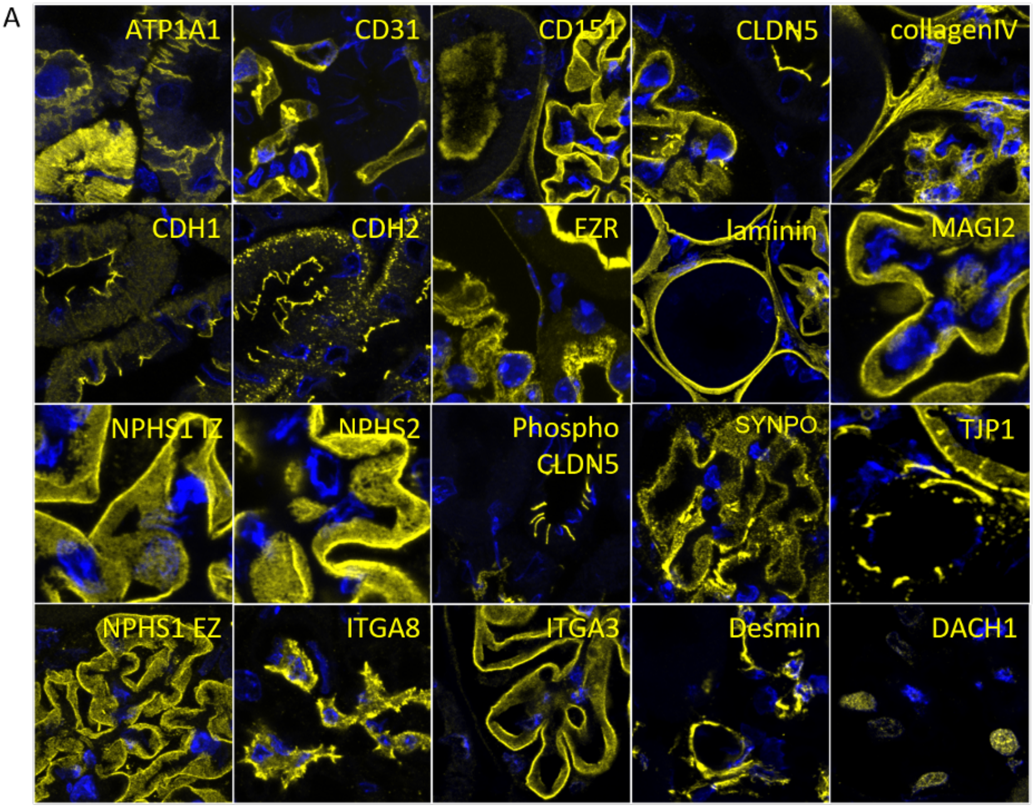


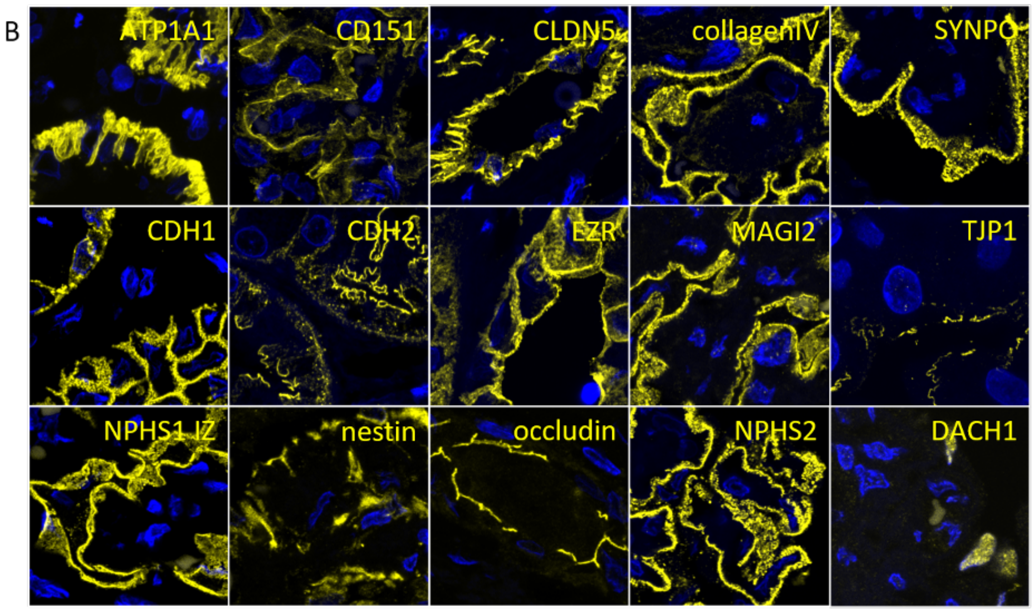


Suppl. Fig. 4 shows single-channel immunofluorescence images from the antibody validation experiment: A total of 20 antibodies targeting murine antigens went into the study (A), while for human targets, 15 antibodies were identified.

Suppl. Fig. 5

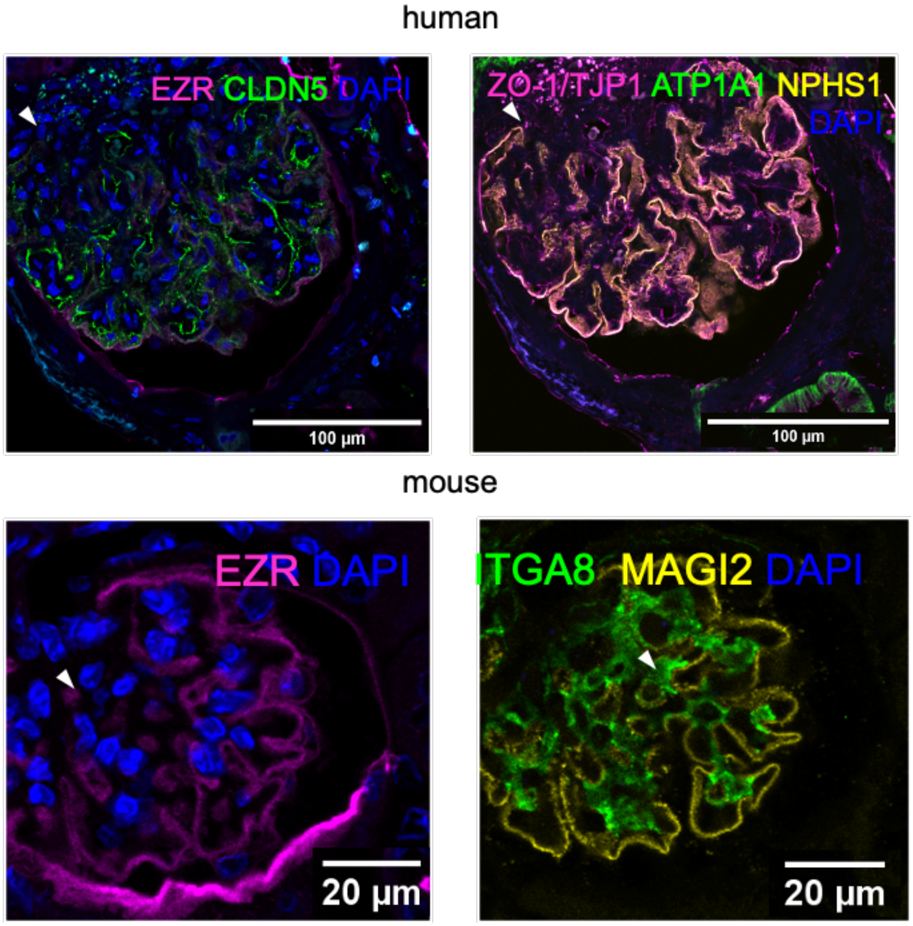


Suppl. Fig. 5 demonstrated inconsistent DAPI staining quality over the imaging rounds independent of the species that was investigated. The scale bars indicate 20 µm.

Suppl. Fig. 6

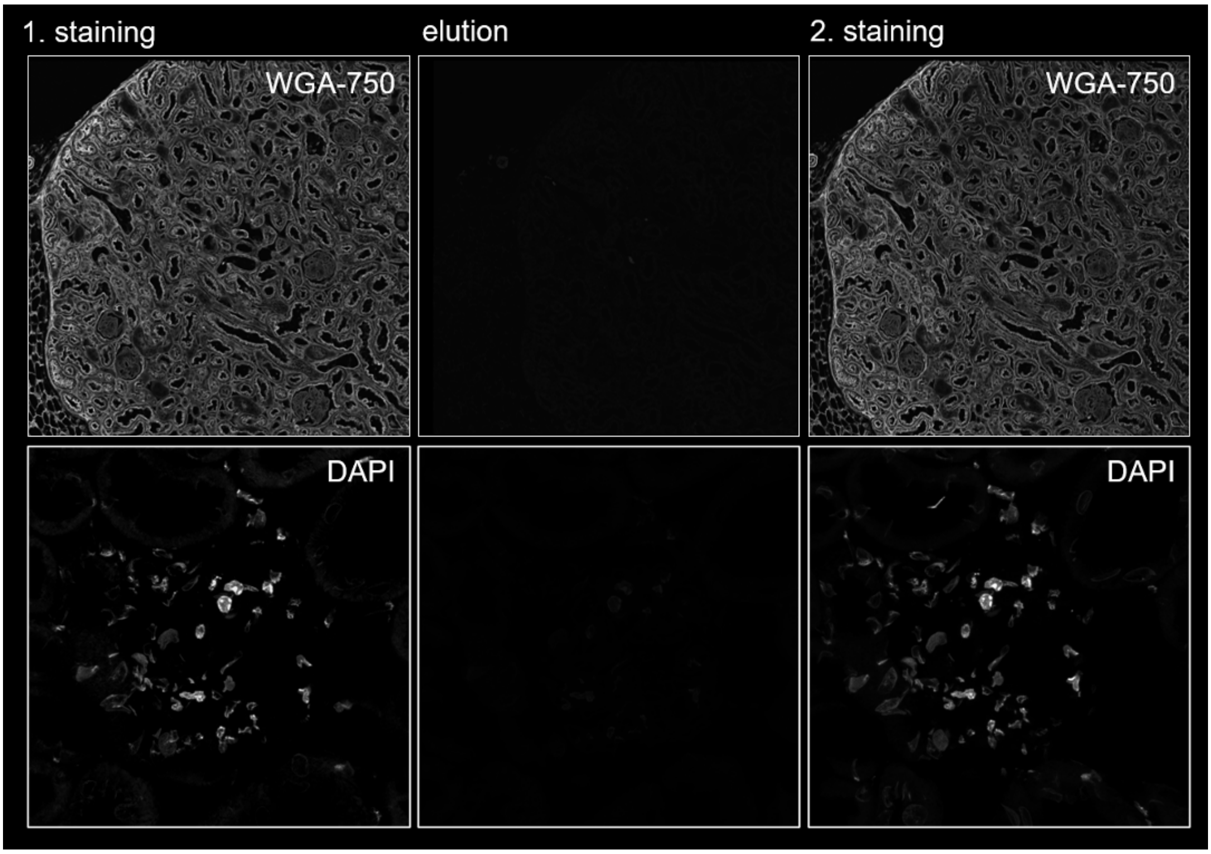


Suppl. Fig. 6: As an alternative marker that could be restained in every round, we identified WGA-770 which could be efficiently eluted and restained on the same section.

Suppl. Fig. 7

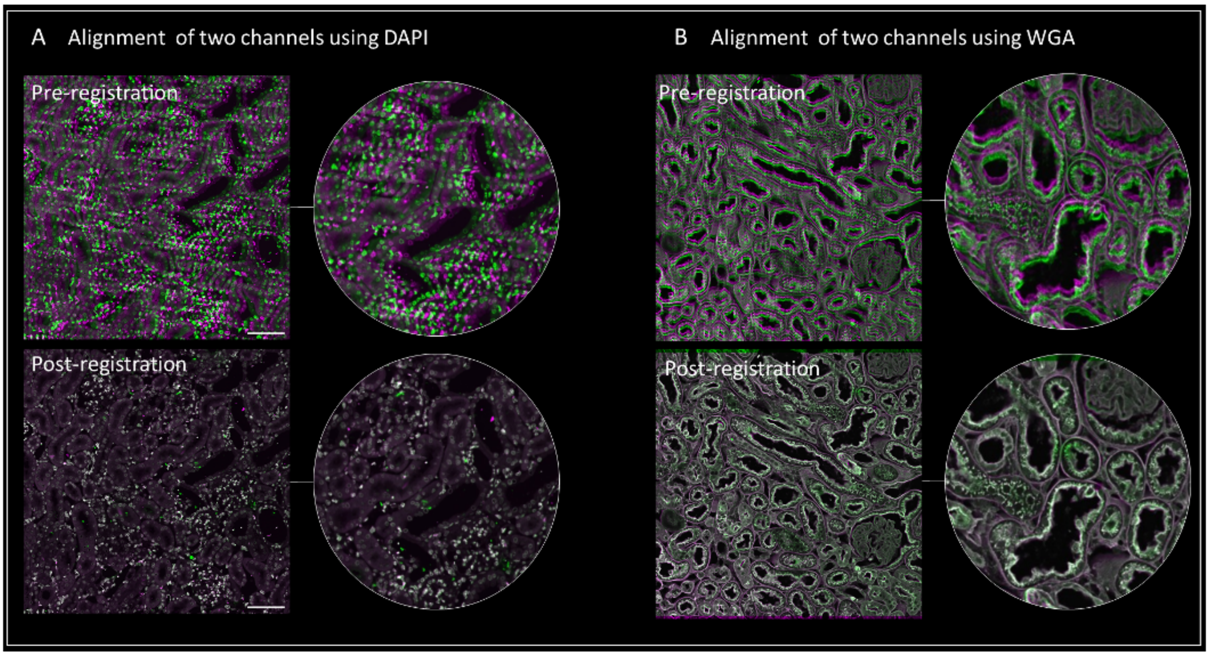


Suppl. Fig. 7 demonstrated the functionality of the shift-registration and correction. Either DAPI-stained cell nuclei, or the WGA staining pattern can be used as a fiducial marker in every staining round. The offset between the imaging rounds is then noticed using the descriptor-based registration in ImageJ and corrected accordingly. Scale bars indicate 100 µm.

Suppl. Fig. 8

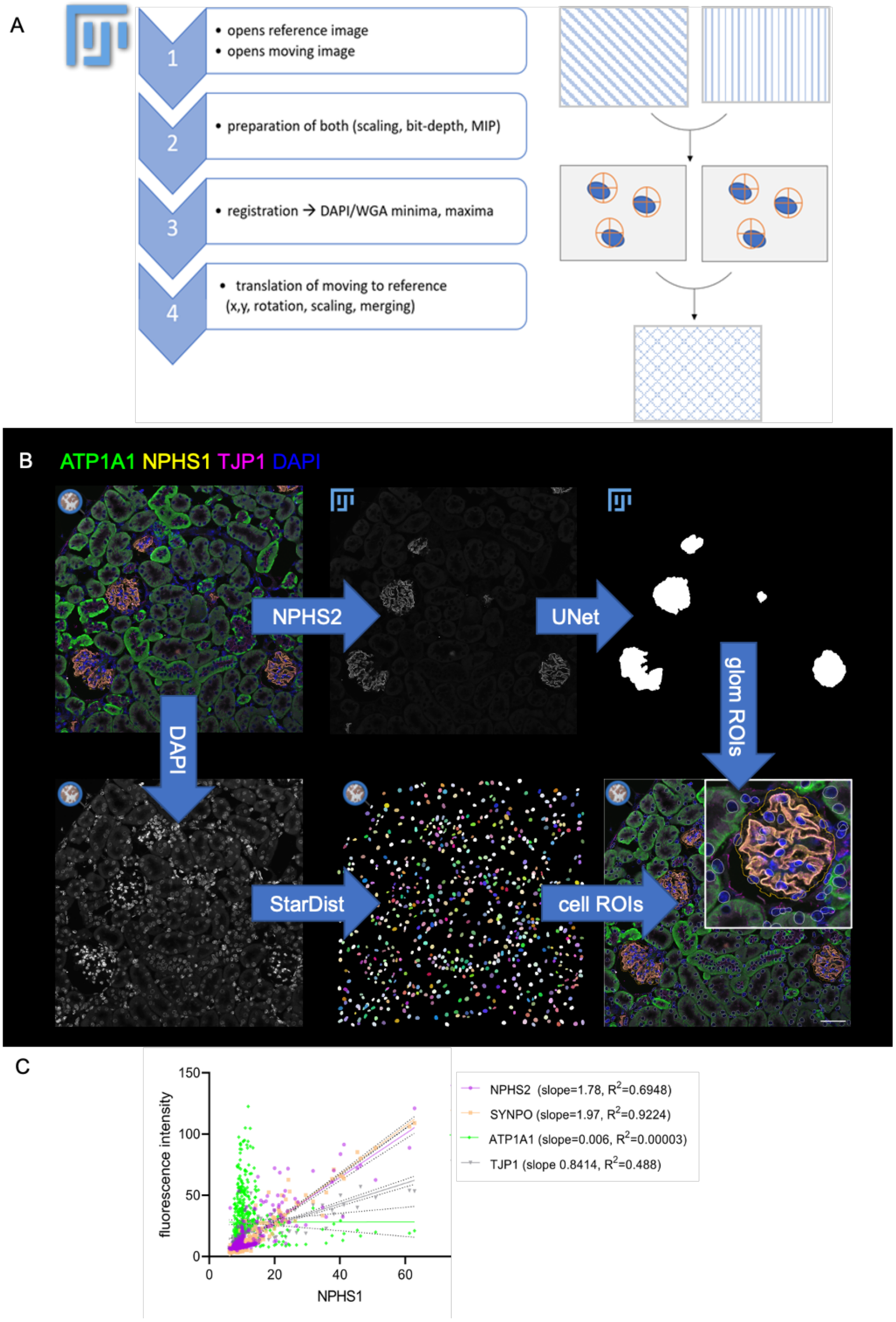


Suppl. Fig. 8 shows the general approach of the quantitative image analysis workflow. First, the image data is registered and aligned (A). B shows the retrieval of single-cell data from the dataset. Glomeruli are segmented using a custom-trained UNet DL-network in FIJI. Single cells are segmented with the StarDist algorithm in QuPath. Finally, the mean fluorescence intensity of all segmented cells is exported (B) enabling single-cell coexpression analysis as shown in B. As expected a positive correlation of the single-cell expression of NPHS and NPHS2, NPHS1 and SYNPO, but not NPHS1 and ATP1A1 could be demonstrated (C).

### Comprehensive material and methods

#### Sialylation of the coverslips

To remove any dust and oily production residuals, the coverslips were cleaned in 96% ethanol on a shaker (100 rpm) for 30 minutes and rinsed twice in distilled water. The completely dried coverslips were incubated with 2% APTES (Thermo Fisher Scientific) in acetone for 30 seconds at room temperature, rinsed in pure acetone, and dried in a laminar flow bench. The coated coverslips were stored dust-free at room temperature for further use.

#### Tissue processing and immunostaining

Mouse tissue was fixed by immersion fixation with 4% paraformaldehyde in PBS pH 7.4 at room temperature overnight. Tissue was embedded in paraffin and 2 µm tissue sections were collected on 22x22mm #1.5 high-precision coverslips coated with 2% (3-Aminopropyl)triethoxysilane (Sigma-Aldrich, Cat. 440140). After deparaffinization, heat-induced epitope retrieval was performed by 5 min boiling in a pressure cooker in TRIS-EDTA buffer pH 9. Coverslips were immobilized in custom-designed and 3D-printed coverslip holders printed with polylactic acid (.stl files available at http://www.github.org/Siegerist). Sections were incubated with 100 mM glycine in PBS for 10 min at room temperature and blocked with 1% BSA, 200 mM $NH_4Cl$, and 150 mM maleimide. Primary antibodies were diluted in 1% BSA; 200 mM $NH_4Cl$ and applied to the sections for 2h at room temperature. Sections were washed with PBS, and secondary antibodies were incubated for 1 h at room temperature. AlexaFluor 555 or CF770-conjugated wheat germ agglutinin (WGA) was added to the secondary antibody solution to a final concentration of 2 µg/ml. After several washes in PBS, cell nuclei were stained with 0.1 mg/ml DAPI. The sections were imaged in 700 mM NAC in $H_2O$ pH 7.4 on a FluoView3000 confocal laser scanning system (Olympus) equipped with a 20x 0.8 NA dry objective (UPLXAPO20X, cyclic immunostainings) or a UPLSAPO60XW NA 1.2 water immersion objective (validation experiments).

#### Antibody elution

After imaging and several washes in PBS, the sections were incubated with 0.5 M L-Glycine; 3 M Urea; 3 M GC; 70 mM TCEP in ddH2O (pH 2.5) for 10 min at room temperature on a shaker (16 rpm). The TCEP was added freshly before each elution. This was repeated 2 times with a final washing step with PBS.

#### Quantitative image analysis

A script (FIJI, http://www.github.org/Siegerist) was created for channel alignment. 2D descriptor-based image registration (registration algorithm Preibisch et al.) was performed using nuclear staining with DAPI or affinity-based carbohydrate staining with wheat germ agglutinin (WGA) as a reference marker for each cycle. Briefly, minima and maxima are detected with sub-pixel accuracy in both respective images and translated to one another using a descriptor-based registration algorithm (Preibisch et al.). This process was iteratively repeated until all images were merged. Next a combined QuPath/Fiji workflow for image classification was set up. Region of interest was marked in QuPath and transferred into the "QuPath ImageJ extension". The ROI was saved as TIFF and a second script (FIJI) was created. It opens the saved TIFF, detects all glomeruli with a pretrained UNet and saves them as ROIs. These are imported in "QuPath ImageJ extension" and sent to QuPath. From the overviews glomeruli were detected with a pretrained UNet, which was used as published and described before. For each cell, the nuclei with surrounding cytoplasm were segmented with a custom-trained Deep Learning network (StarDist) in QuPath (v.0.4.3). This was retrained with … manually segmented nuclei out of DAPI stained multiplexed images.

The dataset was exported from QuPath and saved as a .csv file. UMAPs from this dataset were generated in R Studio using the tidyverse, umap, patchwork, ggplot2, and esquisse packages.
